# Serotonergic treatment normalizes midbrain dopaminergic neuron increase after periaqueductal gray stimulation-induced anticipatory fear in a rat model

**DOI:** 10.1101/714592

**Authors:** Shawn Zheng Kai Tan, Yasin Temel, Ariel Yovela Chan, Andrea Tsz Ching Mok, Jose Angelo Udal Perucho, Arjan Blokland, Luca Acquili, Wei Ling Lim, Lee Wei Lim

## Abstract

**Background:** Electrical stimulation of the dorsolateral periaqueductal gray (dlPAG) in rats has been shown to elicit panic-like behaviour and can be a useful tool for modelling anticipatory fear and agoraphobia.

**Methods:** In this study, we further analysed our previous data on the effects of escitalopram (a selective serotonin reuptake inhibitor, SSRI) and buspirone (a 5-HT1A receptor partial agonist) on dlPAG-induced anticipatory fear behaviour in a rat model using freezing as a measure. We then used tyrosine hydroxylase (TH) immunohistochemistry to probe the effects on dopaminergic neurons.

**Results:** Although acute treatment of escitalopram, but not buspirone, was effective in reducing anticipatory freezing behaviour, chronic administrations of both drugs were comparably effective. We found that the number of dopaminergic neurons in the ventral tegmental area (VTA) was lowered in both chronic buspirone and escitalopram groups. We showed a strong correlation between the number of dopaminergic neurons and freezing in the VTA. We further showed positive correlations between dopaminergic neurons in the VTA and substantia nigra pars compacta in escitalopram and buspirone groups, respectively.

**Limitations:** Although our data strongly hint to a role of dopaminergic mechanisms in the dlPAG induced fear response, more in-depth studies with larger sample sizes are needed to understand the neuronal mechanisms underlying the interactions between serotonergic drugs and dopaminergic cell number and fear behavior.

**Conclusion:** Chronic treatment with an SSRI and a 5-HT1A agonist decrease the number of dopaminergic neurons in the VTA. These effects seem to be associated with reduced dlPAG-induced anticipatory freezing behaviour.

**Key Points:**

- Chronic treatment of escitalopram and buspirone was effective in reducing dlPAG induced anticipatory freezing behaviour.
- The number of dopaminergic neurons in the ventral tegmental area (VTA) was lowered in both chronic buspirone and escitalopram groups and was correlated to freezing.
- We found positive correlations between dopaminergic neurons in the VTA and substantia nigra pars compacta in escitalopram and buspirone groups, respectively.

## Introduction

Electrical stimulation of the dorsolateral periaqueductal gray (dlPAG) in rats has been shown to elicit panic-like behaviours such as vigorous running and jumping in an open-field arena (Lim et al., 2009, 2008). Memory of panic-like behaviour leads to contextual conditioning, where animals associate the context (e.g., the open-field) to fear, which models anticipatory anxiety and agoraphobia (Lim et al., 2010, 2008). Periaqueductal gray (PAG) serotonin transmission has been shown to play a crucial role in regulating the panic response; studies have shown that selective serotonin reuptake inhibitors (SSRIs) can attenuate or increase the threshold for escape behaviours in animals (Hogg et al., 2006; Schenberg et al., 2002; Zanoveli et al., 2005). In this study, we further analysed our previous data (Lim et al., 2010), which studied the effects of escitalopram (an SSRI) and buspirone (a 5-HT1A receptor partial agonist) on dorsolateral PAG(dlPAG)-induced anticipatory anxiety). We used freezing, which is a species-specific defence response defined as the absence of all movement except that required for respiration (Blanchard and Blanchard, 1969), as a measure of fear. It is known that both SSRIs and 5-HT1A receptor agonists exert effects on dopaminergic neurons (Dremencov et al., 2009; Jahanshahi et al., 2010). Further, dopaminergic transmission has been shown to be important for processes and modulation of fear and memory (Abraham et al., 2014; Luo et al., 2018; Pignatelli and Bonci, 2015; Tan et al., 2019a, 2019b). Therefore, we examined the effects of chronic escitalopram/buspirone on dopaminergic neurons by tyrosine hydroxylase (TH)-immunohistochemistry of the ventral tegmental area (VTA), substantia nigra pars compacta (SNpc), dorsal raphe nucleus (DRN), and median raphe nucleus (MnR).

## Results

### Acute administration of escitalopram, but not buspirone, reduced anticipatory freezing behaviour

To test the efficacy of acute BUSP and ESCIT in reducing panic-like freezing behaviour, animals received electrical stimulation or sham stimulation of the dlPAG and were then tested in the open field. At 24 h post-stimulation, animals received either saline, BUSP, or ESCIT and were immediately returned to the open field (Fig 1A). Two-way ANOVA revealed an effect of stimulation, drugs, and their interaction (lowest F = 8.9, all ps< 0.05) on the measured freezing behaviour. Tukey’s posthoc test revealed a significant difference between DBS-SAL and Sham-SAL groups (p < 0.05), indicating DBS induced increased freezing. Tukey’s post-hoc test further revealed a significant difference between DBS-SAL and DBS-ESCIT (p < 0.05), but not between DBS-SAL and DBS-BUSP groups (p = 0.999), which indicated an acute effect of escitalopram, but not buspirone, in reducing anticipatory freezing behaviour (Fig 1C). The heatmap and full table of p-values can be found in Supplementary Figure 1.

**Figure 1.**
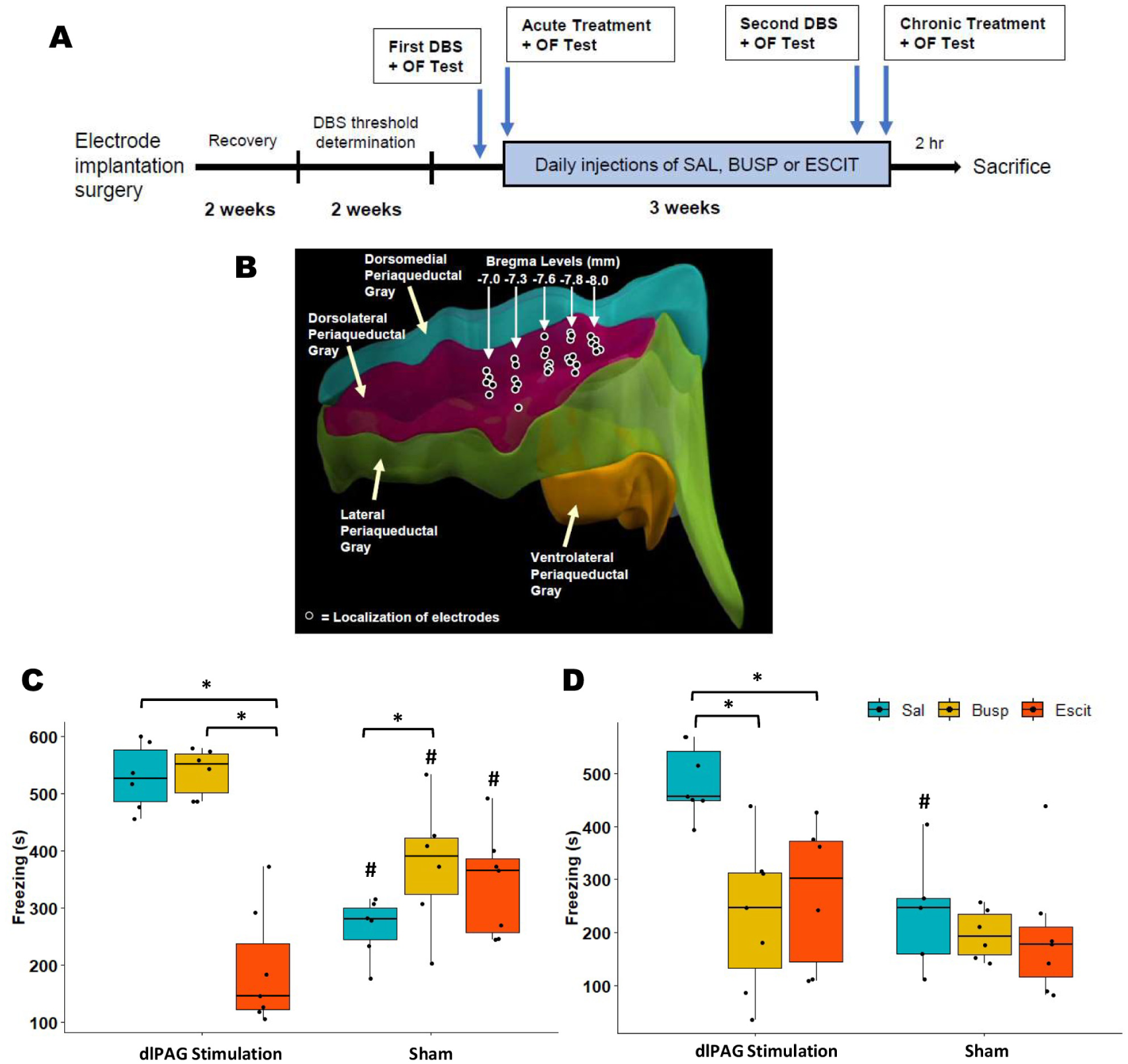
Chronic administrations of buspirone and escitalopram were effective in reducing anticipatory fear freezing behaviours. (A) A schematic representation of the timeline of the experiments. (B) A 3-D reconstruction and lateral view and of the subdivisions of the periaqueductal gray longitudinal columns. The dark dots represent the localization of electrode implantation sites. (C) & (D) show box-plots of the behavioural data from the acute and chronic treatments with saline, buspirone, and escitalopram in the dIPAG stimulation and sham groups, respectively. * represents p < 0.05 between bracketed groups, represents p < 0.05 between corresponding dlPAG stimulation groups.

### Chronic administration of buspirone and escitalopram reduced anticipatory freezing behaviour

To test the efficacy of chronic BUSP and ESCIT in reducing panic-like behaviour, animals received chronic administration of saline, BUSP, or ESCIT for 3 weeks. Animals then received electrical stimulation or sham stimulation of the dlPAG before testing in an open field. At 24 h after stimulation when animals were returned to the open field, two-way ANOVA revealed an effect of drug and DBS (lowest F = 7.7, all ps < 0.05) and a trending interaction effect (F(2,32)=3.2, p = 0.054) on freezing behaviour. Tukey’s post-hoc revealed a significant difference between DBS-SAL and Sham-SAL groups (p < 0.05), indicating DBS induced increased freezing. Tukey’s post-hoc test further revealed a significant difference between DBS-SAL and both DBS-BUSP and DBS-ESCIT groups (all ps < 0.05), but no significant difference between DBS-BUSP and DBS-ESCIT groups (p = 0.98), indicating similar effects of chronic BUSP and ESCIT in reducing anticipatory freezing behaviour (Fig 1D). The heatmap and full table of p-values can be found in Supplementary Figure 1.

### Chronic buspirone and escitalopram treatments and dlPAG DBS caused changes in the number of dopaminergic neurons

To understand the effects of chronic treatment of BUSP and ESCIT and dlPAG DBS on dopaminergic neurons, TH immunohistochemistry and neuronal cell counting were performed on VTA, SNpc, DRN, and MnR brain sections (Fig 2A). Two-way ANOVA revealed an effect of dlPAG DBS on all structures (lowest F = 4.7, all ps < 0.05). There was an effect of drug in VTA (Fig 2B), DRN (Fig 2D), and MnR (Fig 2E) (lowest F = 4.9, all ps < 0.05), but not in SNpc (Fig 2C). There were no interaction effects, although it should be noted that VTA showed a trending interaction effect (F(2,21)= 3.4, p = 0.052). Tukey’s post-hoc test conducted on all structures revealed a significant difference between DBS-SAL and Sham-SAL groups in VTA (p < 0.005), but not in SNpc, DRN, or MnR (all ps > 0.05), which suggests dlPAG DBS induced increased numbers of TH neurons in the VTA. There were a lower number of TH neurons in DBS-BUSP and DBS-ESCIT groups compared to the DBS-SAL group (all ps < 0.01), but no significant difference compared to their sham counterparts, which suggests chronic administration of both BUSP and ESCIT reversed the effects of dlPAG on TH neurons. Tukey’s post-hoc test on SNpc showed no significant differences between all groups (all ps > 0.05), DRN showed significantly lowered TH neuron counts between DBS-SAL and DBS-ESCIT groups (p = 0.02), and MnR showed a significantly lowered TH neuron count between Sham-SAL and Sham-ESCIT (p = 0.04) groups. The heatmap and full table of p-values can be found in Supplementary Figure 1.

**Figure 2.**
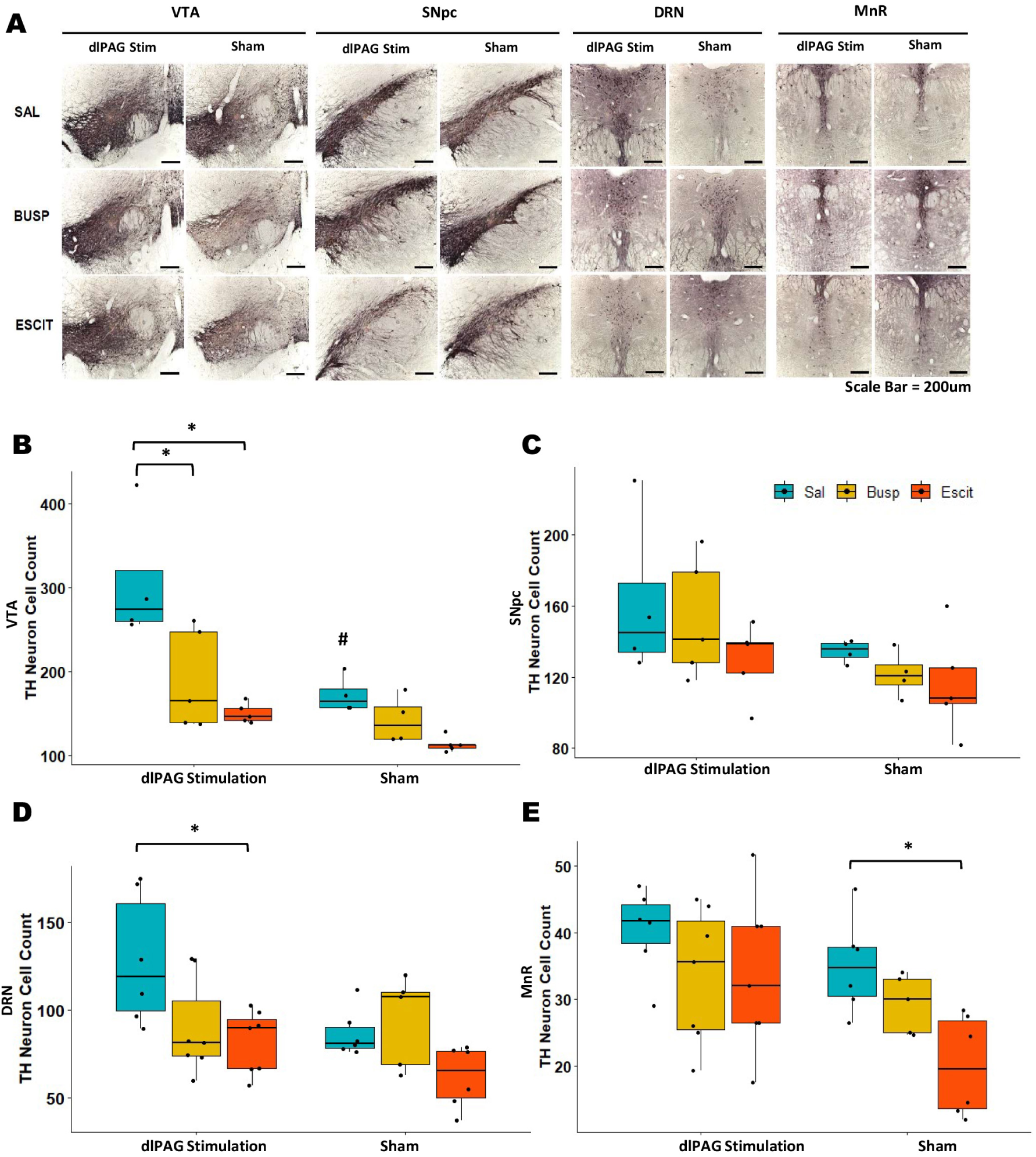
dlPAG DBS and chronic buspirone and escitalopram treatment caused changes in the number of dopaminergic neurons. (A) Representative low-power photomicrographs of VTA and SNpc sections from the brain of rats in dIPAG stimulation and sham groups treated with saline, buspirone, and escitalopram, respectively. Boxplot of TH cell counts for VTA (B), SNpc (C), DRN (D), and MnR (E) showing significantly lowered dopamine neurons counts with dlPAG DBS and chronic buspirone or escitalopram in VTA, and a significantly lowered dopamine neuron counts with escitalopram in DRN. * represents p < 0.05 between bracketed groups, represents p < 0.05 between corresponding dlPAG stimulation groups.

### Escitalopram treatment correlates to TH neuron count in VTA, whereas buspirone treatment correlates to TH neuron count in SNpc

To understand the relationship between chronic administration of BUSP or ESCIT and dopaminergic neurons, individual drug groups and TH cell counts in VTA, SNpc, DRN, and MnR were analysed by Spearman’s rank correlation (Fig 3B, D, F, H, respectively). ESCIT showed significant positive correlation with TH neuron count in the VTA (R = 0.68, p = 0.06), whereas BUSP showed significant positive correlation with TH neuron count in SNpc (R = 0.95, p < 0.001). No significance was seen in DRN or MnR for either drug. All R and p values are shown in the respective graphs (Fig 3B, D, F, H).

**Figure 3.**
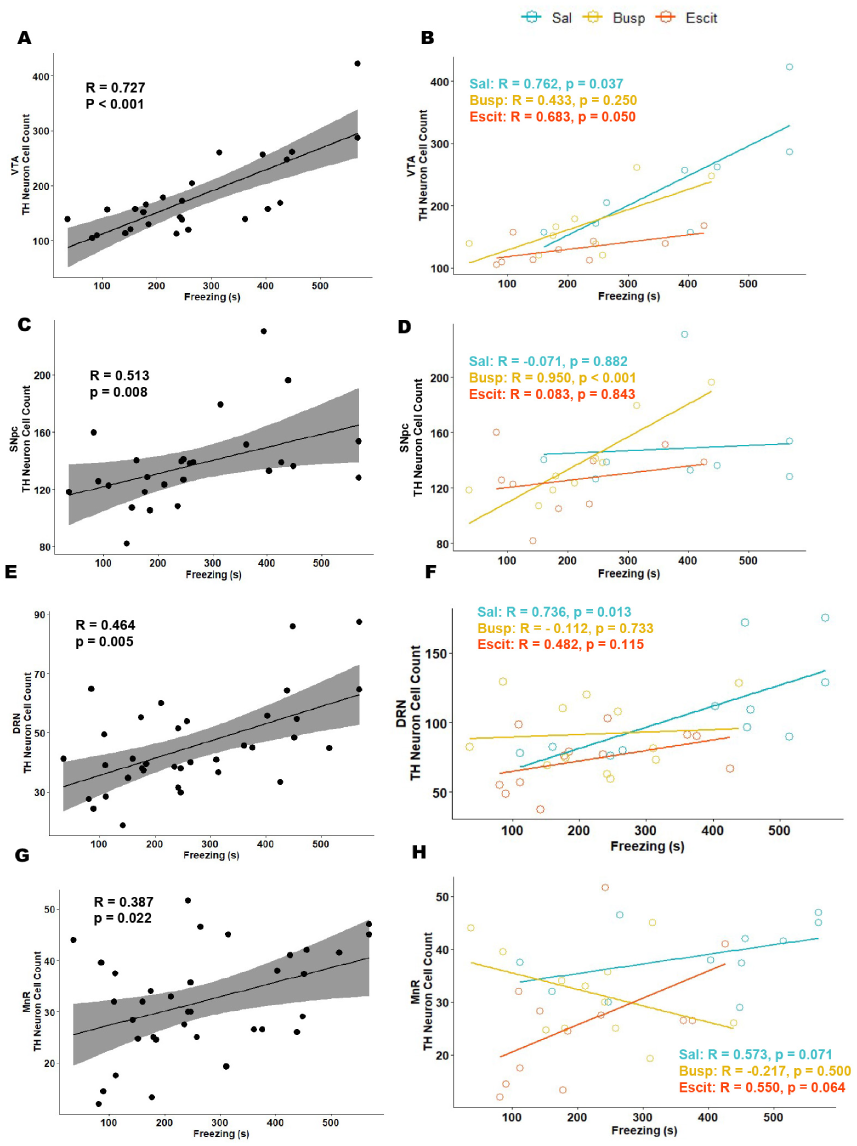
Correlation plots of TH neuron cell count and freezing behaviour Global correlation plots showed significant positive correlations between TH neuron count and anticipatory fear freezing behaviour in VTA (A), SNpc (C), DRN (E), and MnR (G). In individual drug groups, escitalopram showed a significant correlation in VTA (B), and a significant correlation in SNpc (D). No significant correlation of buspirone or escitalopram was seen in DRN (F) or MnR (H).

## Discussion

In this study, we extended our previous findings to show that acute treatment of ESCIT, but not BUSP, was effective in reducing anticipatory freezing behaviour (mimicking agoraphobia). However, both drugs were comparably effective in reducing freezing behaviour after chronic administration. We further demonstrated that chronic BUSP or ESCIT decreased the number of dopaminergic neurons in the VTA, whereas only chronic ESCIT decreased dopaminergic neurons in the DRN. Lastly, we showed that there was a strong positive correlation between the number of dopaminergic neurons and freezing in the VTA, and positive correlations between dopaminergic neurons with ESCIT and BUSP treatments in the VTA and SNpc, respectively. This suggests that both drugs, while both reduce the number of dopaminergic neurons, exert their effects via different dopaminergic-related neuronal circuits.

The use of dlPAG stimulation is advantageous in that it directly induces the activation of the panic-circuit independent of the behavioural context and is highly reproducible (Hogg et al., 2006; Jenck et al., 1995; Lim et al., 2011). This feature is useful as an unconditioned stimulus for modelling agoraphobia. The use of freezing to measure fear response (rather than distance moved or corner time) is more relevant in cases where there is a lack of shelter or no close predators (Eilam, 2005). Our behavioural data agree with previous studies showing anxiolytic drugs (including serotoninergic drugs) were effective in reducing panic-like responses in dlPAG stimulation models (Hogg et al., 2006; Jenck et al., 1998, 1995, 1990). This study extended our previous findings (Lim et al., 2010) and showed that ESCIT and BUSP reduced freezing behaviour.

The mesolimbic dopamine pathway appears to play an important role in fear learning and memory, including encoding and consolidation (Pezze and Feldon, 2004; Pignatelli and Bonci, 2015). Therefore, it is not surprising that freezing activity was correlated to dopaminergic structures (namely VTA, SNpc, DRN, and MnR) in our model. Furthermore, both BUSP and ESCIT have been shown to affect dopaminergic transmission. BUSP was shown to increase circulating dopamine (Lechin et al., 1998) and occupy dopamine receptors (Ciano et al., 2017). ESCIT was shown to increase the firing rate of VTA dopamine neurons (Ivanov and Konradsson-geuken, 2011), although it should be noted that another study found an opposite effect (Dremencov et al., 2009). How serotonin modulators affect dopaminergic systems is still largely unknown and more studies are needed to fully understand their effects. Interestingly, the individual drug correlations appear to point towards ESCIT exerting its effects through/on VTA dopaminergic neurons, whereas BUSP exerts effects through/on SNpc dopaminergic neurons. This is in contrast to the TH cell count data, which showed both drugs exerted their effects mainly through/on the VTA. Nevertheless, BUSP has been shown to have an anti-dyskinetic effect on a Parkinson’s disease model (Azkona et al., 2014) and the SNpc is suggested to play a crucial role in this effect (Sagarduy et al., 2016; Sharifi et al., 2012), which implies that BUSP directly affects the SNpc, how much this affects anticipatory anxiety requires further mechanistic work. We should note that a limitation of this study is the relatively small sample size in terms of biological replicates (although multiple technical replicates for cell counts on each sample was performed, with averages of them being used for analysis), which might influence correlation studies. This has also limited our ability to obtain meaningful data from correlation studies breaking down dlPAG DBS and sham groups – though freezing as a measure would be sufficient given dlPAG DBS function was to induce fear which is measured by freezing. Overall, while the data suggest a complex interplay between VTA and SNpc dopaminergic neurons in the observed effects, more investigation is needed to fully understand this relationship.

## Conclusions

In this study, we showed that both chronic administration of BUSP and ESCT were effective in reducing anticipatory fear, though likely mediated via different structures and pathways in the dopaminergic system. A major limitation has been the lack of mechanistic studies on how drugs affect dopaminergic systems, as well as the inconsistencies in the literature. More work is needed to fully understand the link between serotonergic drugs and their effects on dopaminergic neurons in fear behaviour.

## Methods

### Animals

Adult male Wistar rats (n=40, 12 weeks old) were individually housed in standard cages with food and water available ad libitum. The environmental conditions were maintained at a temperature of (21.1 °C) and humidity (60-65%) humidity in a reverse 12 h/12 h light/dark cycle. This study was approved by the Animal Experiments and Ethics Committee of Maastricht University, as well as the Committee on the Use of Live Animals in Teaching and Research (CULATR) of The University of Hong Kong.

### Experimental groups

Rats were randomly divided into six experimental groups: three groups receiving dlPAG deep brain stimulation (DBS) and three groups receiving sham stimulation and treated with either escitalopram (DBS-ESCIT, n=7; SHAM-ESCIT, n=7), buspirone (DBS-BUSP, n=7; SHAM-BUSP, n=6), or saline (DBS-SAL, n=7; SHAM-SAL, n=6), respectively. A schematic representation of the timeline of the dlPAG DBS and drug treatments is shown in Figure 1 (A).

### Surgical procedures

The surgical procedures were performed as previously described (Lim et al., 2010; Temel et al., 2007). Rats were anaesthetized during the entire procedure using a combination of ketamine (90 mg/kg) and xylazine (10 mg/kg) injected subcutaneously. Rats were placed in a stereotaxic frame (Stoeling, Wood Dale, USA) and electrodes were implanted at the level of the dlPAG (from Bregma: anterposterior, −7.6 mm; mediolateral, +0.7 mm; ventral, −4.8 mm; coronal approach angle of 10º)(Paxinos and Watson, 2006) and secured to the skull using dental cement. The electrodes consisted of gold-plated needle combined with an inner wire of a platinum-iridium (Technomed, Beek, the Netherlands; IDEE Instruments, Maastricht University) with a tip and shaft diameter of 50 µm and 250 µm, respectively. All rats received a subcutaneous injection of Temgesic (0.1 mg/kg) for pain relief directly after the surgery and were allowed to recover for 2 weeks.

### DBS procedures

The DBS procedures for inducing fear-like behaviour in rats were performed according to previous studies (Lim et al., 2010, 2009). To determine the level of the escape threshold, the dl-PAG DBS groups had a preliminary session in their home cages. The stimulation amplitude was gradually increased until escape behaviour was observed. At each step, the stimulation period was 15 s followed by a stimulation-off period of 45 s. The stimulation frequency was set at 50 Hz and pulse width at 0.1 ms based on our previous studies (Lim et al., 2010, 2009). The stimulation was delivered using a World Precision Instruments (WPI) digital stimulator (DS8000, WPI, Berlin, Germany) and a stimulus isolator (DLS100, WPI, Berlin, Germany). Real-time verification of the DBS stimulation parameters was monitored using a digital oscilloscope (Agilent 54622D oscilloscope, Agilent Technologies, Amstelveen). Animals requiring stimulation intensities above 100 µA were excluded from the analysis. After the threshold determination session, all rats were allowed a recovery period of 2 weeks. Sham animals were similarly connected to the stimulator, but no current was delivered.

### Drug administration

The dosage and administration of BUSP and ESCIT in animals have been shown to be effective in our previous study (Lim et al., 2010). A week prior to the actual experimental tests, all animals received 1 mL saline injections three times on alternating days to habituate them to the injection procedure. Animals were administered a single acute dose of drug before the first open-field behavioural test. Escitalopram oxalate (ESCIT; H. Lundbeck A/S, Copenhagen, Denmark) and Buspirone hydrochloride (BUSP; TOCRIS, Cookson Inc., Missouri, USA) were dissolved in saline (SAL; 0.9% NaCl). The animals received a subcutaneous injection (in the volume of 1 mL/kg) of ESCIT (10 mg/kg) 60 min before or BUSP (3 mg/kg) 120 min before the first open-field behavioural test. For testing the chronic effects of these drugs, animals received daily injections of ESCIT (10 mg/kg), BUSP (3 mg/kg) or SAL for 21 days. A final dose of SAL, ESCIT and BUSP was administered (at 60 min and 120 min, respectively) before the second open-field behavioural testing to investigate the chronic treatment effects.

### Behavioural evaluation

Freezing behaviour of rats was evaluated in an open-field (square: 100 cm × 100 cm; height: 40 cm) made of clear plexiglass with an open top and a dark floor (Lim et al., 2010). The rat was placed in the open-field arena for 10 min and their behaviour was recorded using a camera. Freezing is defined as the absence of all movement except that required for respiration (Blanchard and Blanchard, 1969). The freezing response was manually scored by researchers who were blinded to the experimental design and animal groups.

### Histological processing

At 2 h after the final behavioural test, rats were anaesthetized with Nembutal (75 mg/kg) and then perfused transcardially with Tyrode (0.1 M) and a fixative solution containing paraformaldehyde, picric acid and glutaraldehyde in phosphate buffer (pH 7.6). Rats were post-fixed for 2h and incubated overnight in sucrose solution, and subsequently frozen rapidly with CO_2_ and stored at −80 oC. Brain tissues were sequentially cut on a cryostat into 30 μm thick sections in the coronal plane and stored at −80 oC. One series of the brain sections were processed for TH immunohistochemistry. Briefly, brain sections were incubated overnight with mouse anti-TH antibody (1:100, kindly provided by Dr C. Cuello, Canada) diluted in 0.1% bovine serum albumin (BSA) and Tris-Buffered Solution (TBS)-Triton (TBST) solution, followed by biotinylated donkey anti-mouse IgG (diluted 1:400, Jackson Immunoresearch Laboratories Inc, Westgove, USA). Sections were then incubated with avidinbiotin-peroxidase complex (diluted 1:800, Vectastain Elite ABC kit, Burlingame, USA) for 2 h. Between steps, sections were washed with TBS and TBST. Tissue sections were incubated with 3, 3’-diaminobenzidine tetrahydrochloride/nickel chloride solution to visualise the horseradish peroxidase (HRP) immune complex. The reaction was stopped after 10 min by washing the sections thoroughly with TBS. Subsequently, all sections were mounted on the Superfrost micro-slides (VWM, Illinois, USA) and cover-slipped with Permount mounting medium (Thermo Fisher Scientific, Waltham, USA). Another series of brain sections were stained with haematoxylin-eosin (Merck, Darmstadt, Germany) to evaluate the localisation of the electrode implantation sites (Fig 1B).

### Evaluation of the TH-immunoreactive cells

Cell counts of TH-immunoreactive (TH-ir) cells were performed in the VTA, SNpc, DRN and MnR brain sections from the DBS and SHAM treatment groups. For quantification of TH-ir cells, three or four sections of the VTA, SNpc, DRN and MnR per animal were observed under an Axiophat 2 imaging microscope (Carl Zeiss Microscopy GmbH, Gottingen, Germany) by an investigator who was blinded to the treatment groups. Photomicrographs of TH-ir cells within the areas of interest were captured using an Olympus DP73 digital camera (Olympus, Hamburg, Germany) attached to a bright-field microscope. Brightness and contrast of the photomicrographs were adjusted in Adobe Photoshop (Adobe Systems, San Jose, USA).

### Statistics

Statistical analysis was conducted in R (version 3.5.2) and visualizations were performed using the “ggplot2” package (Wickham, 2016). As linearity of data cannot be assumed, outliers (n=2; 1 DBS-SAL, 1 DBS-BUSP) in the behaviour data were removed using the ROUT method (Motulsky and Brown, 2006). No outliers were removed from the cell count and correlation studies.

## Acknowledgements & Disclosures

This research was funded by a Hong Kong RGC-ECS Grant (27104616) that awarded to L.W.L.. W.L.L. was the recipient of the International Brain Research Organization-Asia Pacific Regional Committee (IBRO-APRC) Exchange Fellowship to work on this project. All authors declared no biomedical financial interests or potential conflicts of interests.

