## Supplementary Figure 1 for "Serotonergic treatment normalizes midbrain dopaminergic neuron increase after periaqueductal gray stimulation-induced anticipatory fear in a rat model"

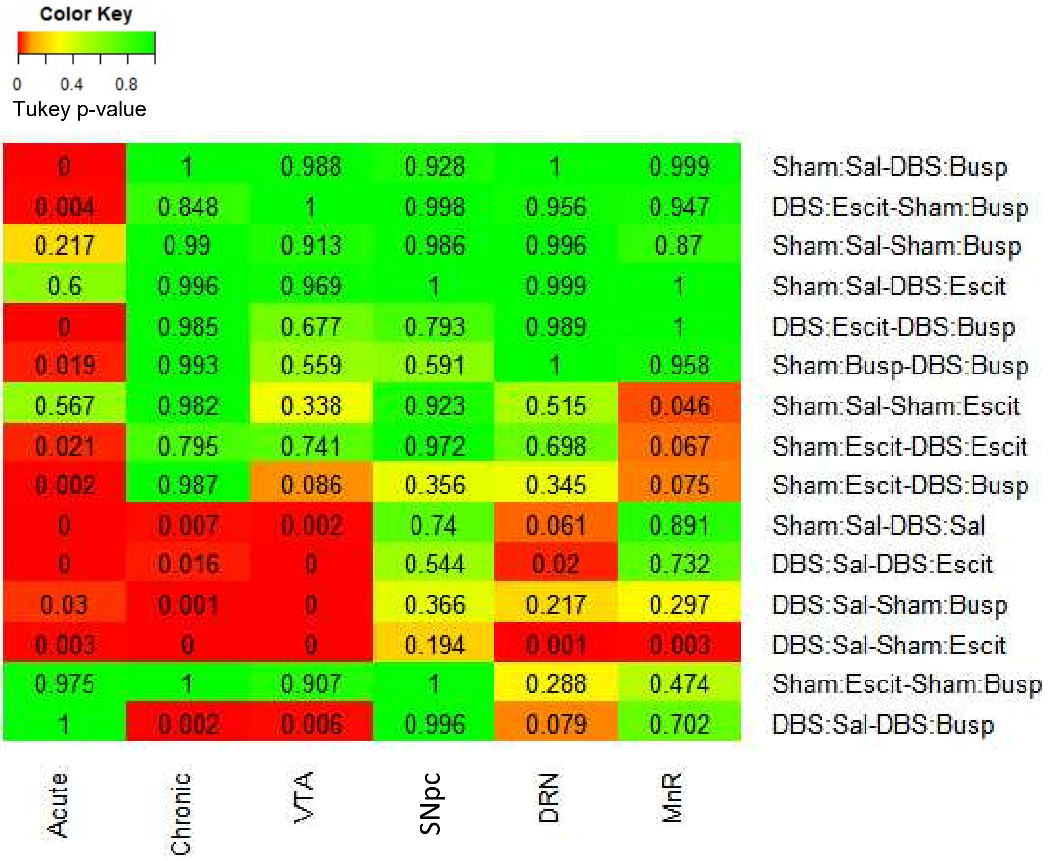


**Supplementary Figure 1. Heat plot of p-values from Tukey's test.** Numbers in the table are p-values from Tukey's post-hoc tests from the ANOVAs. The heatmap shows the levels of significance (red represents more significance and green represents less significance).
